# TransCRISPR - sgRNA design tool for CRISPR/Cas9 experiments targeting DNA sequence motifs

**DOI:** 10.1101/2022.04.05.487109

**Authors:** Tomasz Woźniak, Weronika Sura, Marta Kazimierska, Marta Elżbieta Kasprzyk, Marta Podralska, Agnieszka Dzikiewicz-Krawczyk

**Author notes:** correspondence; Strzeszyńska 32, 60-479 Poznań, Poland.

## Abstract

Eukaryotic genomes contain several types of recurrent DNA motifs, e.g. transcription factor motifs, miRNA binding sites, repetitive elements. CRISPR/Cas9 can facilitate identification and study of crucial DNA motifs. We present transCRISPR, the first online tool dedicated to search for DNA sequence motifs in the user-provided genomic regions and design optimal sgRNAs targeting them. Users can obtain sgRNAs for chosen DNA motifs, for up to tens of thousands of target regions in 30 genomes, either for the Cas9 or dCas9 system. TransCRISPR provides user-friendly tables and visualizations, summarizing features of identified motifs and designed sgRNAs such as genomic localization, quality scores, closest transcription start sites, and others. Experimental validation of sgRNAs for MYC binding sites designed with transCRISPR confirmed efficient disruption of the targeted motifs and effect on expression of MYC-regulated genes. TransCRISPR is available from https://transcrispr.igcz.poznan.pl/transcrispr/

## INTRODUCTION

Several types of recurrent DNA motifs exist in eukaryotic genomes and are important components of complex regulatory networks^1^. For example, transcription factors (TFs) recognize specific sequence motifs in DNA and regulate target gene expression^2^. MiRNAs bind to their target transcripts via short complementary seed sequences^3^. Splicing also depends on the recognition of specific sequences by the spliceosome^4^. The role and significance of these DNA motifs might be examined by their blocking or disruption, e. g. with use of CRISPR/Cas9 system^5–10^. Clustered Regularly Interspaced Short Palindromic Repeats/Cas9 (CRISPR/Cas9) has become one of the most powerful tools for genome editing and revolutionized genome engineering. However, to perform reliable and informative CRISPR/Cas9 experiments, high specificity and efficiency of the approach are required. In answer to this need, numerous online tools have been designed. Several online CRISPR/Cas9 tools are available which allow designing the most optimal single-guide RNAs (sgRNAs) targeting specific sequences and predict their off- and on-target scores to increase their specificity and efficiency^11−14^. Although they offer a wide range of possibilities, none of them allows searching for a specific DNA motif in a sequence and designing sgRNAs targeting this motif.

Here, we present a highly versatile online tool, transCRISPR, created to identify DNA motifs in the sequence of interest and to design sgRNAs with optimal off- and on-target scores. It can be applied both for single sequences as well as for large lists of genome coordinates, enabling design of sgRNA libraries for genome-wide CRISPR/Cas9 screens.

## RESULTS

TransCRISPR is an online tool dedicated to designing sgRNAs targeting various DNA motifs. This software performs several steps to calculate and display data for given input: 1) processing sequences; 2) processing motifs and finding motifs in sequences; 3) calculating off-targets and on-targets; 4) finding localization and calculating statistics.

### User interface

To run a query, the user has to provide input data and select available options (Figure 1). In Step 1, the reference genome is selected. Currently, one can operate on thirty genome assemblies, including e.g. human, mouse, rat, fruit fly, zebrafish, *C. elegans* and others. We plan to further expand the list of available genomes in near future.

**Figure 1.**
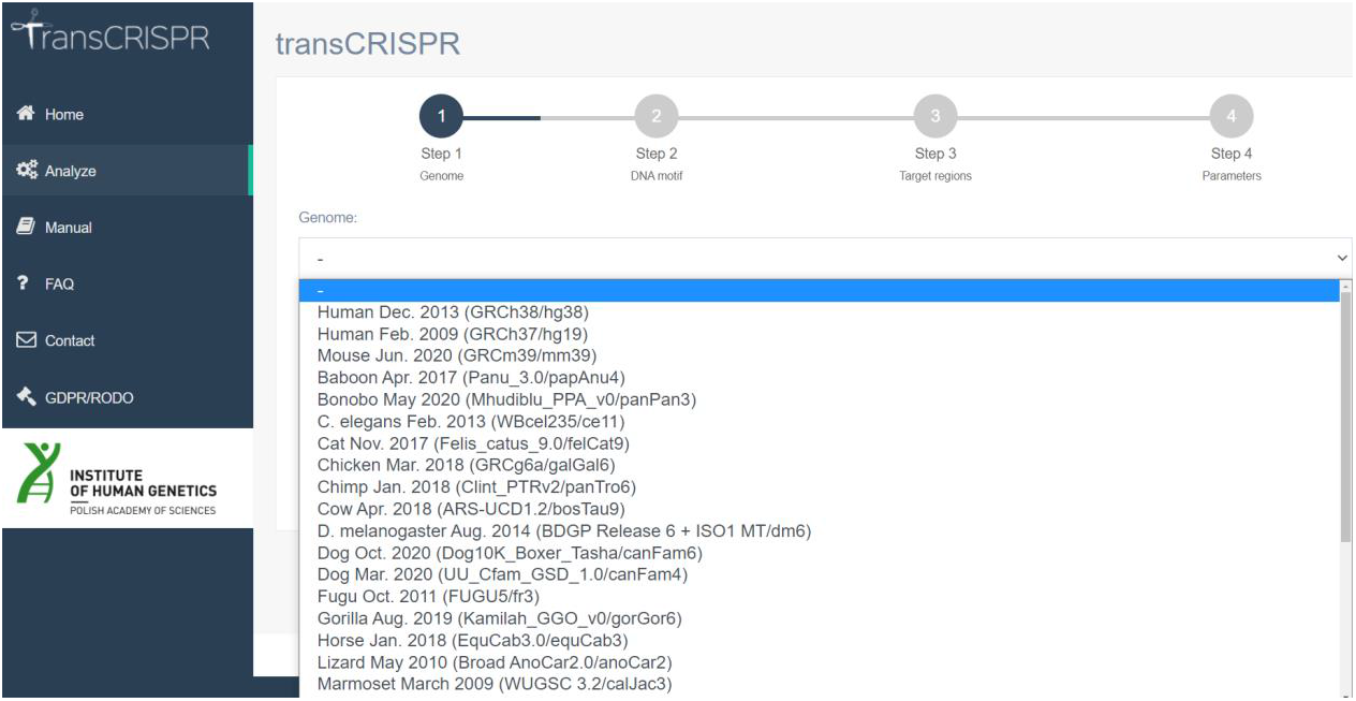
TransCRISPR interface and query steps.

TransCRISPR offers many ways of submitting queries. In Step 2 DNA motifs might be entered directly in the window as FASTA or comma separated format, or as a motif matrix in various formats, including output files of programs analyzing transcription factors (e. g. JASPAR, TRANSFAC). The motifs might be also uploaded as a sequence or matrix file. Motifs provided as a sequence can contain A, C, G, T nucleotides or IUPAC codes. Next, in Step 3 the target regions where DNA motifs will be searched for are provided. Target sequence may be pasted directly in the window as a text (in FASTA or coma separated format) or as genomic coordinates; those can be uploaded as files as well. In the last step, one may choose from various modes of motifs and guides search.

If the motif sequences were entered directly in the form of “Motif sequences” or in the form of “Motif sequence file”, they will be all searched separately. However, if they were entered in the form of a motif matrix (“Motif matrix” or “Motif matrix file”), the user can choose criteria according to which motifs will be generated from the matrix. The first and recommended option is to use a set of D. R. Cavener rules for consensus sequence^15^. The nucleotide having over 50% frequency at the specific position and simultaneously having the frequency more than twice higher than the second most common nucleotide is regarded as the consensus nucleotide. If this criterium is not met, two nucleotides whose sum of the frequencies exceeds 75% are regarded as consensus nucleotides; if none of these criteria is met, N is assigned to the position. Alternatively, the user can set that motifs will be generated including all nucleotides which cover at least x% (5%, 10%, 15%, 20%, 25%) at a given position, or the most frequent nucleotides which together cover at least 80%, 85% or 90% at the given position in the motif.

Next, one can choose between the *S. pyogenes* Cas9 and dCas9 variants which define how the sgRNAs are searched with respect to the motifs. In the Cas9 variant, only those guides that lead to the cut within the DNA motifs (taking into account that the cut occurs 3 nt upstream of PAM) are designed. In the dCas9 variant, any guides that overlap with at least one nucleotide of DNA motifs are included. As a standard, for all found sgRNAs off-targets up to 4 mismatches are analyzed. To reduce the analysis time, the “rapid” option can be chosen which includes only off-targets with up to 3 mismatches. The user can optionally enter their e-mail address to be informed when calculations are done. Moreover, during the analysis, the user is informed about its progress and current step, as well as about the queue of tasks. For convenience, the details of the query might be checked later (Figure 2). Results are available on the website for seven days.

**Figure 2.**
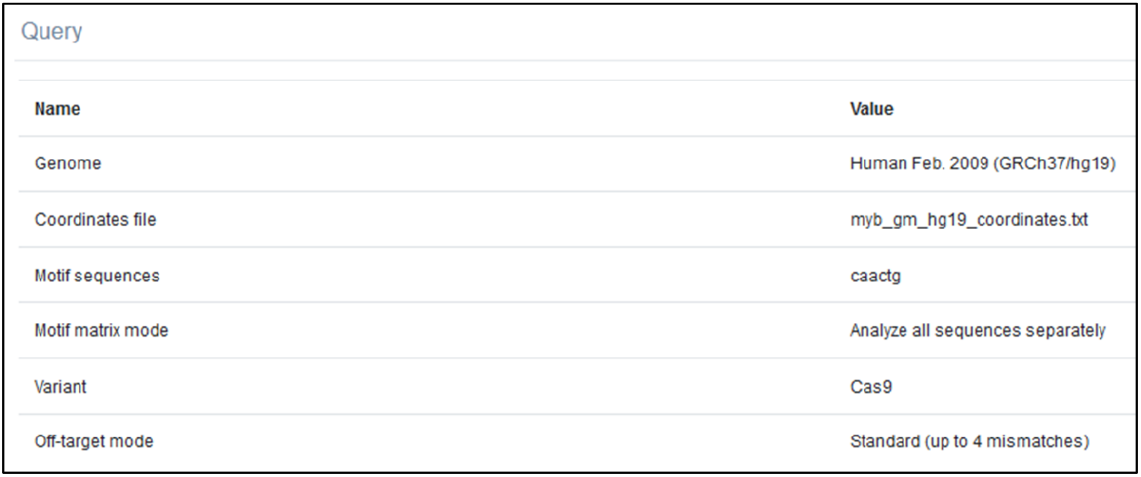
Summary of query details available on the result page.

### Analysis results and data visualization

On the results page, the user is first informed about the number of found motifs and sgRNAs, an average number of guides per motif, and average on- and off-target scores, which are additionally presented on histograms. Distribution of guides per motif and their genomic localization are presented on pie charts. These charts can be viewed full screen, printed or downloaded in various formats. Information about found motifs includes: motif sequence, their position in the uploaded genomic sequence and localization in relevance to genes (whether it is in coding or non-coding exon, intron or between genes), including the genomic positions of transcription start sites (TSSs) for the closest up- and downstream gene. Next, sgRNAs found for the motifs are presented in the table, where their sequence, relative position in the sequence and DNA strand are shown, together with the calculated on- and off-target scores. It is possible to view the details of the most significant off-targets. Information about the genomic localization of motifs and closest TSSs is not available if target regions were provided as sequences; genomic coordinates are required to obtain the full characterization.

The results may be additionally filtered according to several parameters. The user may choose to include only motifs within a specific genomic localization. While working with TF motifs, for Cas9 option, we recommend to include only motifs in non-coding exons, introns and intergenic, since cutting within motifs located in coding exons will likely disrupt the protein and hence the results will not be conclusive for the TF binding. For dCas9 option, we recommend excluding motifs localized in the window −200 nt/+100 nt relative to TSS as this region is the most effective for CRISPR interference. Thus, it would be difficult to determine whether the observed effect is a result of blocking the TF binding or of interfering with gene transcription due to dCas9-induced mechanisms. sgRNAs can be filtered based on the on- and off-target threshold. The user may a priori remove sgRNAs having off-targets with 0 or 1 mismatches, regardless of their CFD score. We recommend that the cut-off of 30 is used for CFD off-target score. It is also possible to filter out motifs for which no guides could be designed.

In case several sgRNAs are found per motif, the user may wish to include only a given number of the best guides. For this purpose, the maximum number of sgRNAs based on the off- or on-target score can be requested. The order in which the guides are displayed can be also changed (default by on-target score; off-target or position are possible). In some instances, DNA motifs may overlap partially and in such a situation some sgRNAs may be duplicated in the motif view. To obtain the list of nonredundant sgRNAs, the user should switch to the “Unique guides” mode. In this view, the information about the sequence and the position of sgRNAs together with on- and off-target scores and targeted motifs are provided.

The user may download the analysis results in several formats (xslx, csv, tsv, bed). In the Excel file, separate sheets provide results per motifs or per unique guides. It is also possible to download the track to visualize motifs and guides together with their on- and off-target scores coded by colors in the UCSC Genome Browser. Moreover, by clicking Display in Genome Browser the user is directly taken to the Genome Browser with this track loaded.

A detailed explanation of preparing the query and analyzing results is provided in the manual available on the transCRISPR webpage, also as a downloadable pdf. To get familiar with transCRISPR and available options, users are advised to run one of the preloaded examples.

Summary of the features available in transCRISPR and comparison with other available tools for sgRNA design is presented in Table 1.

**Table 1.**
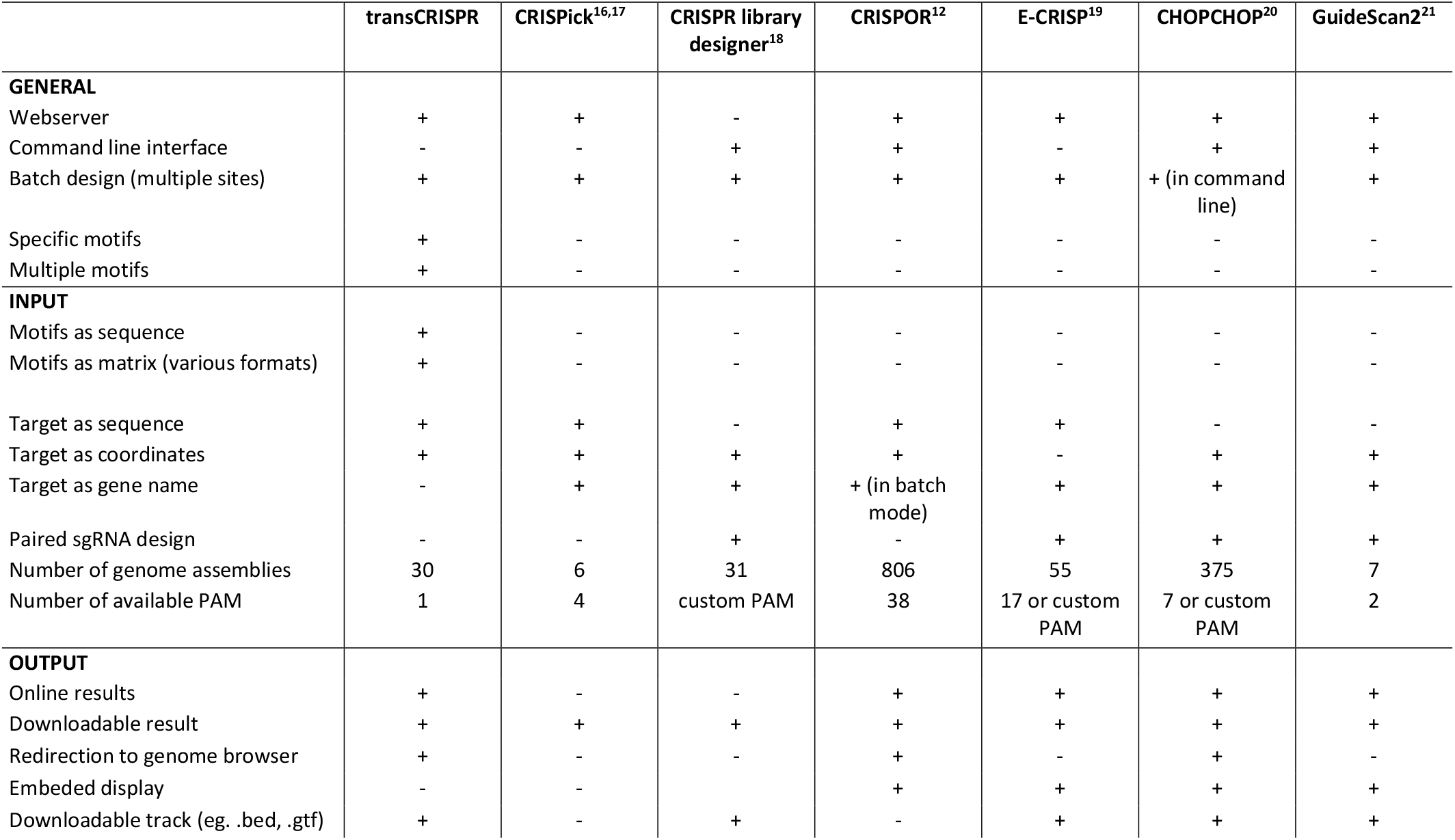
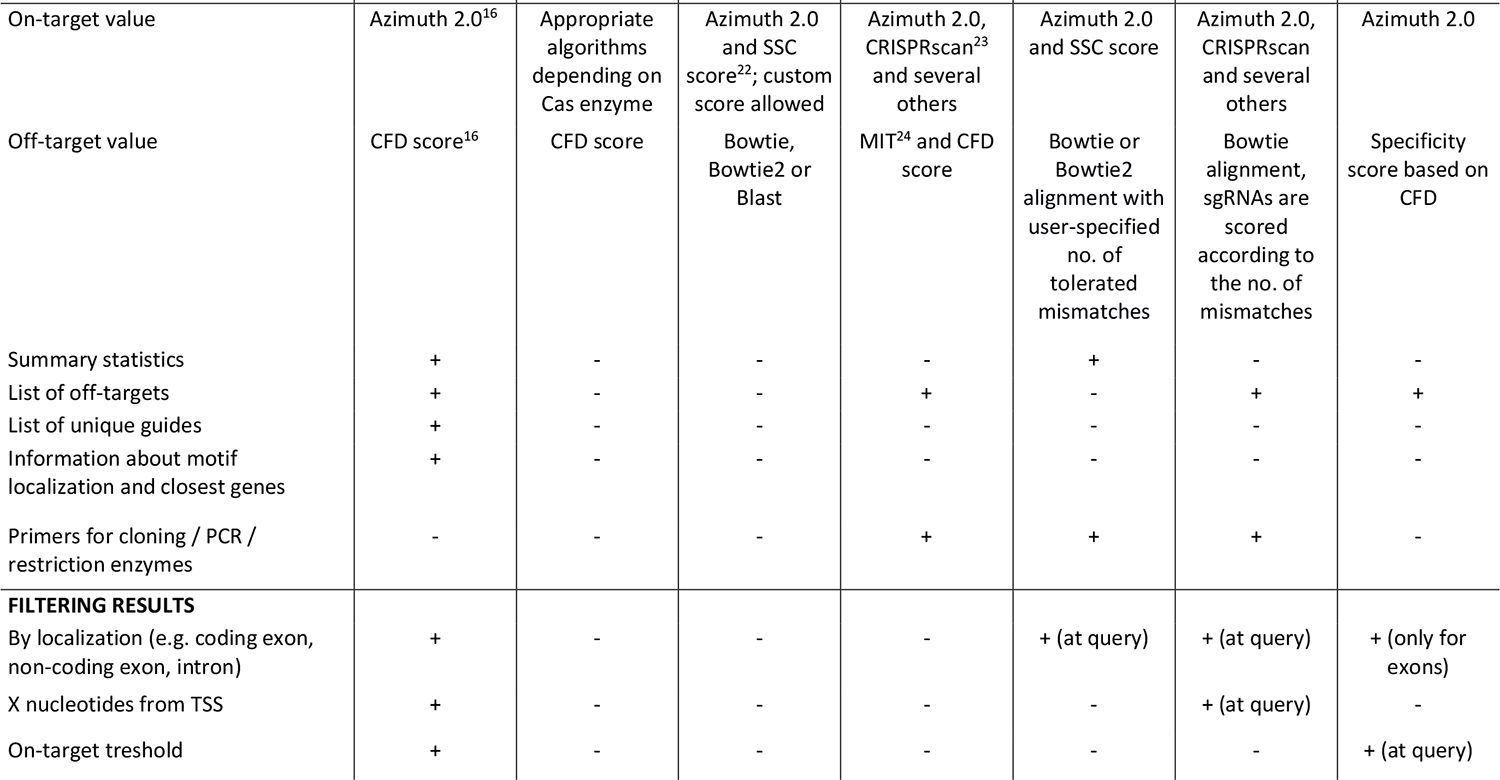
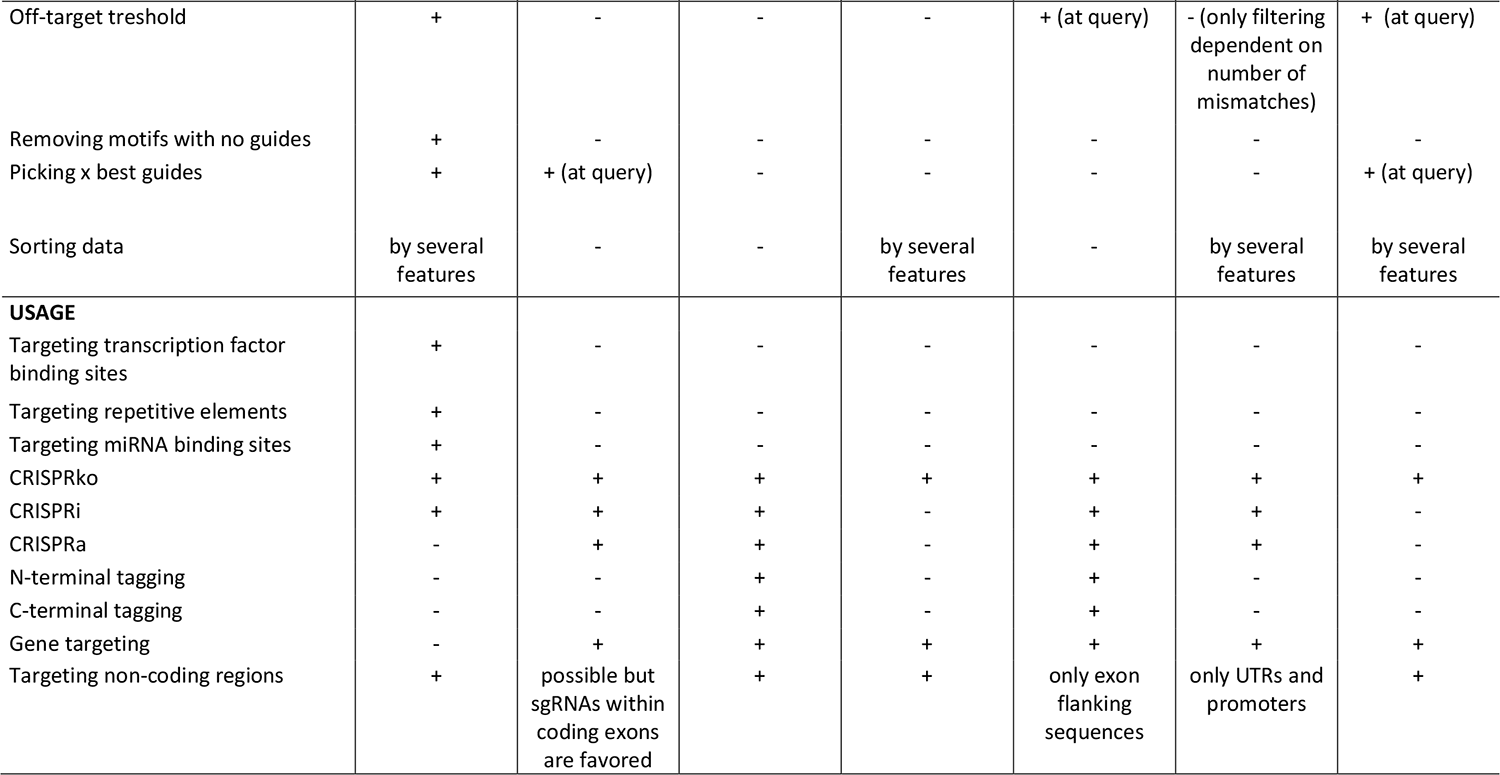
Comparison of features offered by transCRISPR and other sgRNA design programs.

### Example of library design

We used transCRISPR to find motifs and sgRNAs within the ChIP peaks for the MYB transcription factor in human GM12878 cells. For this purpose, we chose GRCh37/hg19 genome assembly and pasted the most common MYB motif sequence (CAACTG) directly into the motif window. Next, we used the coordinates of MYB ChIP peaks retrieved from UCSC Table Browser, track ENCODE 3 TFBS, table GM12878 MYB, and uploaded them as a coordinates file (3748 peaks). First, we chose Cas9 variant and standard off-target mode (up to 4 mismatches) (Figure 2).

After the calculations are finished, the upper panel shows the summary of motifs and guides (Figure 3A)– there are 975 found motifs and 947 guides targeting them, which gives 0.97 guides per motif. The detailed information about the number of sgRNAs per motif presented on the pie chart below shows that for 40.7% of the motifs no sgRNAs could be designed. The remaining motifs were targeted mostly by 1 or 2 sgRNAs; for some up to 6 sgRNAs were designed. The average on- and off-target scores are above 50 which indicates an overall good quality of designed sgRNAs. Detailed information provided on histograms shows that the majority of sgRNAs have off-target scores above 70, only a few are below 50. The predicted cutting efficiency is medium as the majority of sgRNAs have the on-target score around 50. Looking at the genomic localization of identified motifs, we observe that the vast majority is localized in the introns or non-coding exons, much less in intergenic regions and only a few in coding exons.

**Figure 3.**
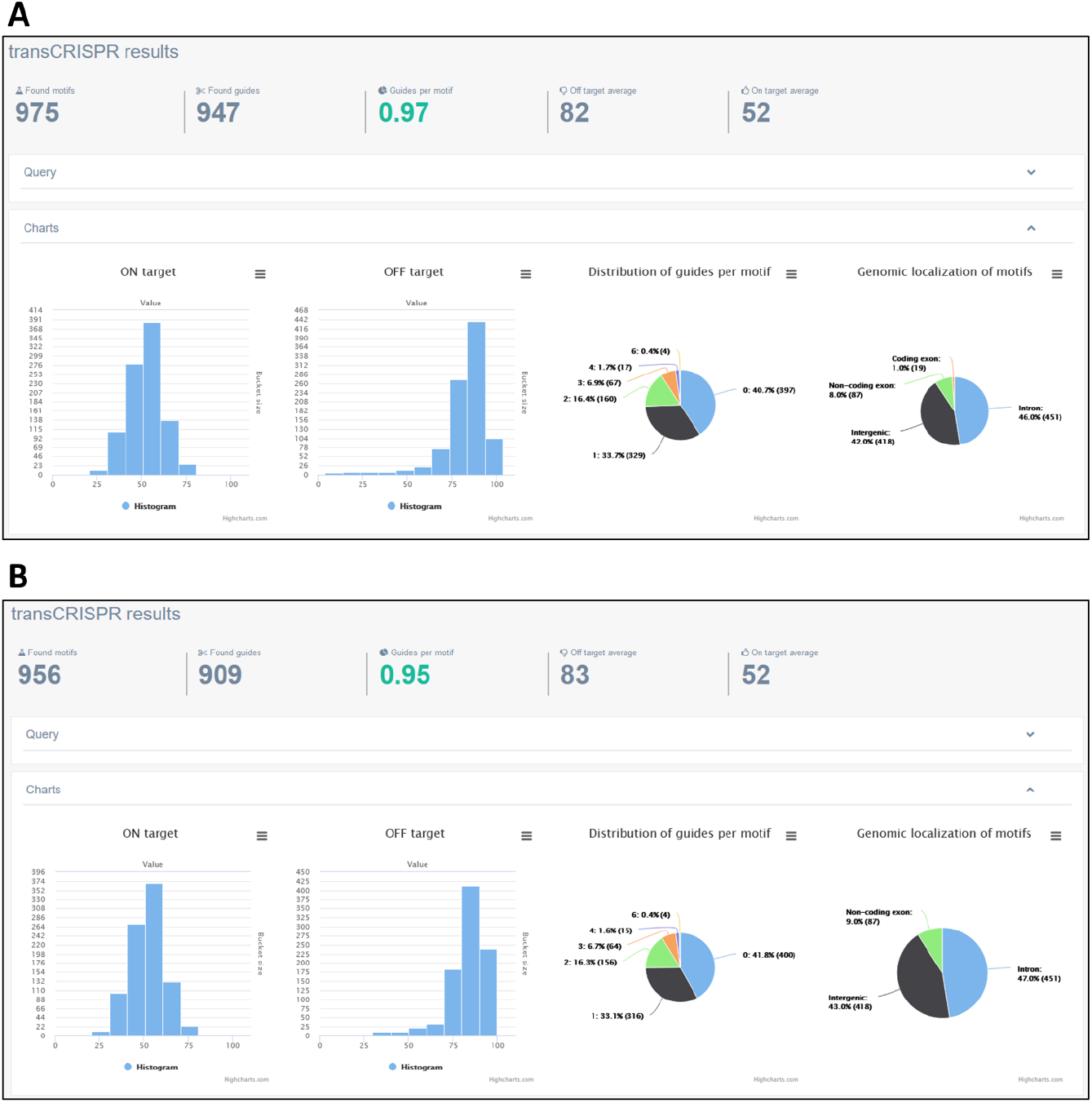
Statistics of transCRISPR results for the example query in Cas9 mode. The screenshots present initial results of analysis described in the example (A) and results after filtering by genomic localization of motifs and off-target scores (B).

As mentioned previously, results can be filtered based on various criteria. We used this option to exclude motifs present in coding regions and guides presenting off-target scores below 30. As a result, we obtained 956 motifs and 909 guides (0.95 guides per motif) with the average off-target score increased to 83. We can see on charts that indeed now there are no motifs in coding exons and no guides with off-target score below 30 (Figure 3B). Figure 4 shows various modes of presenting results by transCRISPR – they can be displayed as a list at the result page (Figure 4A), as a list downloaded in xslx format (Figure 4B and C) or visualized in Genome Browser (Figure 4D). Colors of sgRNAs bars depict on- and off-target values range.

**Figure 4.**
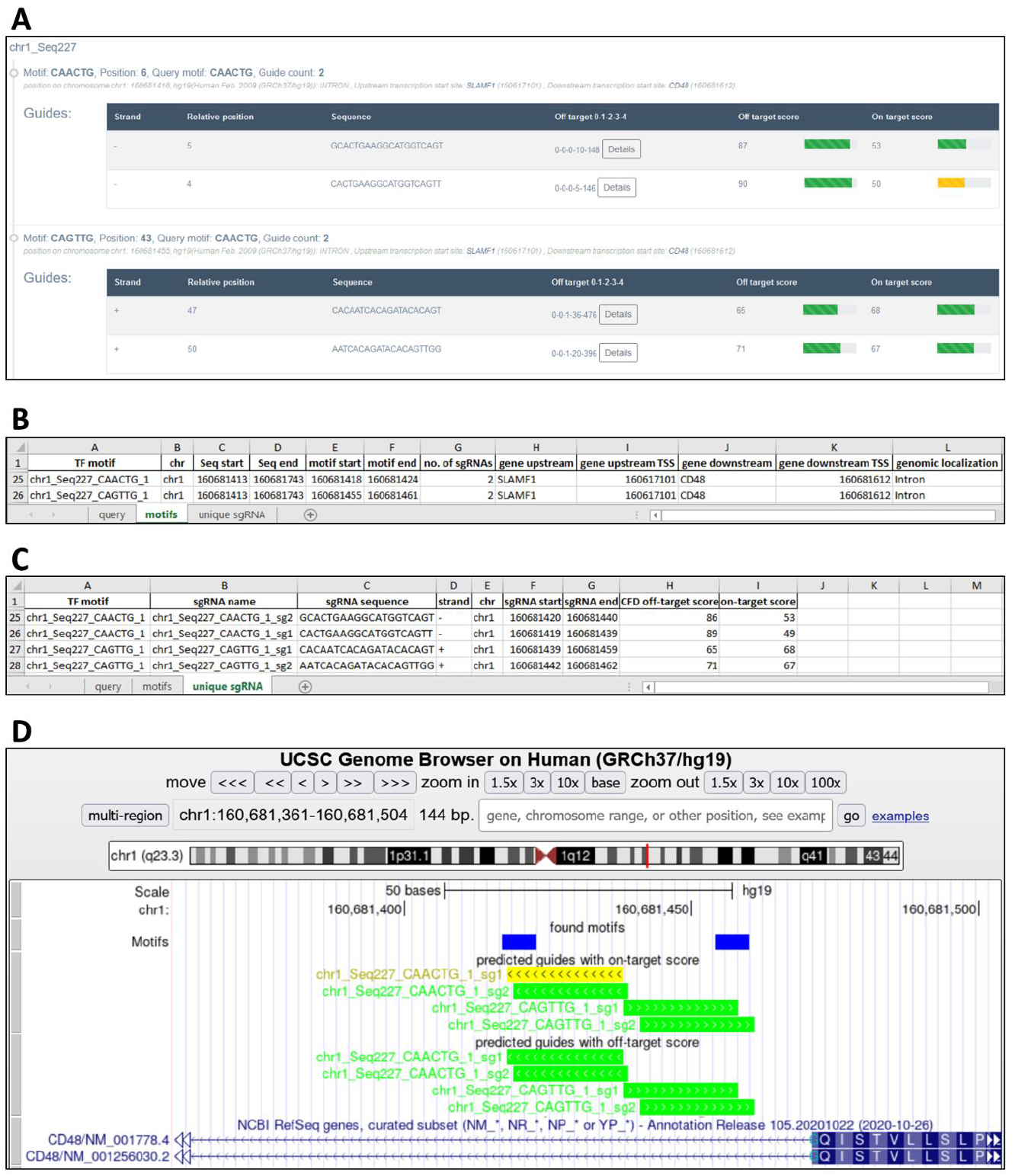
Different modes of viewing transCRISPR results. The screenshots present the same motifs and guides (A) directly on the results page; (B) and (C) in the downloaded xslx file (information on motifs – B, information on unique guides - C); and (D) as tracks in Genome Browser.

Changing the Cas9 variant in the query to dCas9 yielded 3285 guides (3.37 guides per motif) (Figure 5A). This is expected, as the rules for sgRNA design are broader in this option. In line with this, the number of guides per motif is more diversified (up to 8), with the prevalence of 1-4 sgRNAs per motif. Average on- and off-target values as well as their distribution on histograms look quite similar to the ones from Cas9 mode. When the filters recommended for dCas9 mode were applied, i.e. motifs localized between −200 nt and +100 nt relative to TSS were excluded, the number of found motifs went down to 873 and the number of designed guides decreased to 2815. Other statistics changed accordingly (Figure 5B).

**Figure 5.**
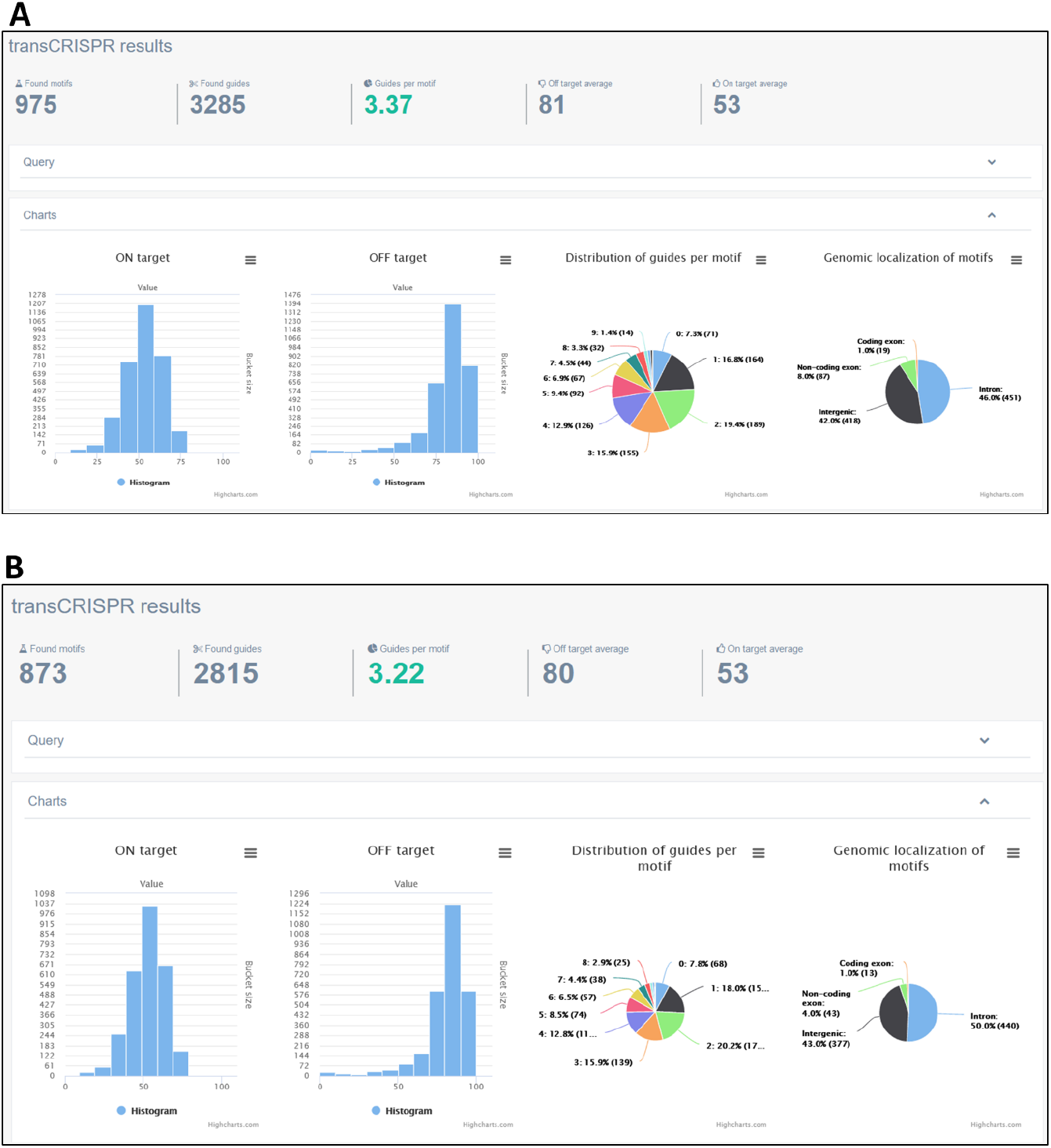
Statistics of transCRISPR results for the example query in dCas9 mode. The screenshots present initial results of analysis described in the example (A) and results after filtering by localization of motifs relative to TSS (B).

### Experimental validation of transCRISPR design

To confirm that transCRISPR is able to design sgRNAs efficiently targeting DNA sequence motifs, we took as an example genes involved the purine biosynthesis pathway which are known to be regulated by MYC: *PPAT, GART, PFAS, PAICS* and *ATIC*. MYC-ChIP peaks proximal to these genes were retrieved from ENCODE data for K562 cells and used as the target sequence in transCRISPR, while the MYC motif matrix was obtained from Jaspar. For each MYC peak (*PPAT* and *PAICS* are localized in a head-to-head orientation with the MYC peak within their promoter) transCRISPR identified 1-3 MYC binding motifs and designed sgRNAs targeting them (Figure 6A-D). sgRNAs showed good specificity (75-99) and moderate predicted efficiency (52-75). Designed sgRNAs were cloned into the lentiCRISPRv2_puro vector and used to transduce K562 cells. Based on TIDE analysis, all sgRNAs resulted in efficient DNA editing (100%, only for ATIC E-box 2 – 78%) and disruption of the E-box motifs (Figure 6A-D, Supplementary Figure 1). Importantly, CRISPR editing of E-box sequences decreased expression of the studied genes. The most pronounced effect was observed for PFAS. Disruption of the intergenic E-box localized between PPAT and PAICS did not affect either gene, while targeting the intronic E-boxes within PAICS reduced expression of PAICS but not PPAT (Figure 6E).

**Figure 6.**
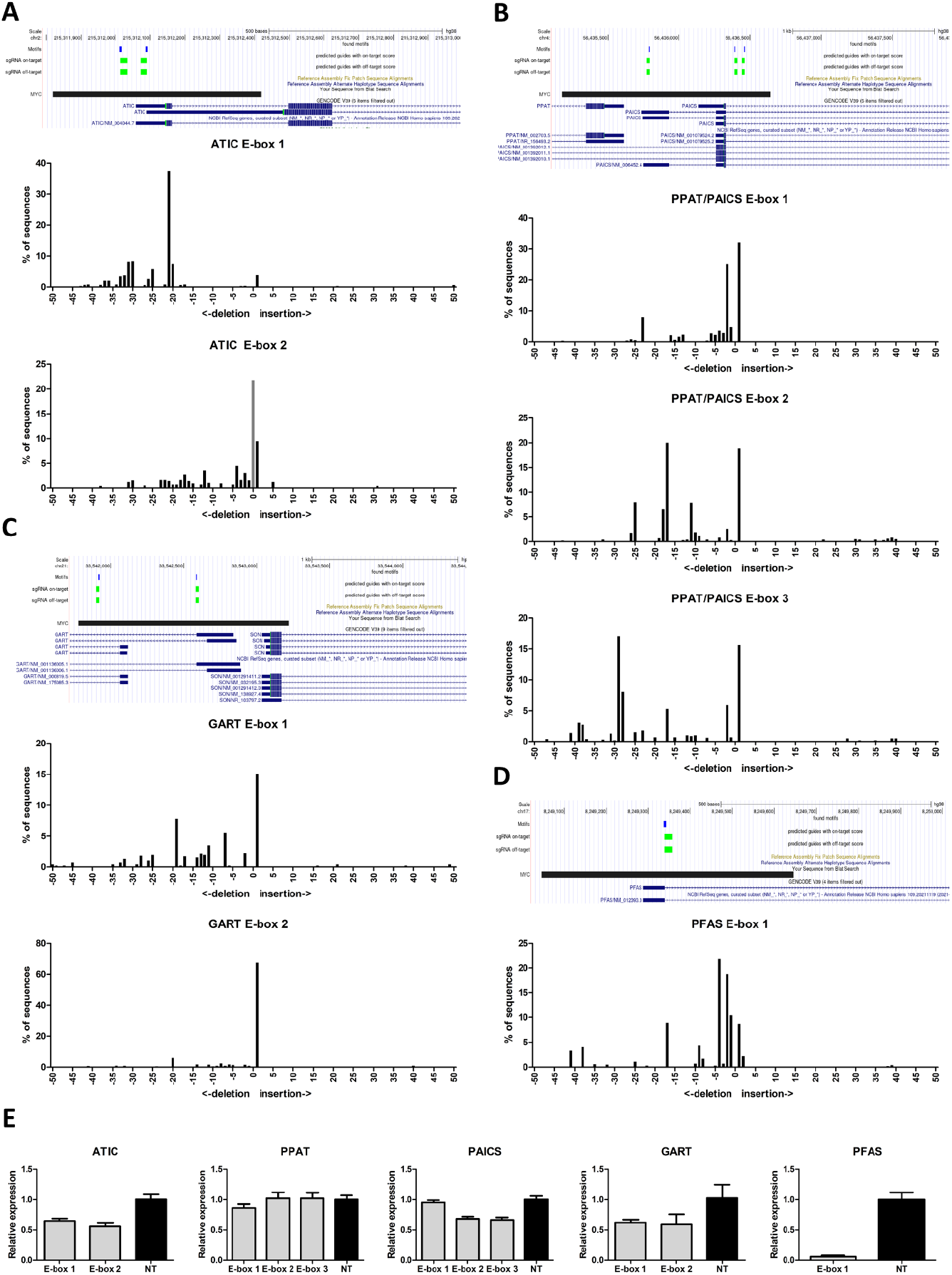
Experimental validation of sgRNAs designed by transCRISPR. A-D) Genomic localization of identified E-box motifs (blue boxes) and targeting sgRNAs (green boxes) within the MYC-ChIP peaks (black boxes). Column charts present results of TIDE analysis for sgRNAs targeting each E-box. E) Expression of genes localized near targeted E-boxes compared to control non-targeting (NT) sgRNAs (average of two NT constructs). Shown are the means and standard deviations from two independent experiments, each in triplicate.

This experiment demonstrated that transCRISPR enables design of efficient sgRNAs for targeted sequence motif disruption, which allows designing and performing experiments to answer biologically relevant questions.

## DISCUSSION

We designed transCRISPR to facilitate the study of DNA motifs. This is a unique tool that enables design of sgRNAs targeting a particular sequence in the region of interest and provides an important novel functionality for sgRNA design algorithms. The ability to disrupt or block sequence motifs can significantly facilitate research and understanding of their function. Importantly, transCRISPR offers wide range of functionalities and options to tailor the results to the user’s needs and is applicable for small queries as well as design of genome-wide sgRNA libraries.

The biggest improvement introduced by transCRISPR is a versatile, efficient and well tested pipeline, which includes both custom code and available tools and algorithms. This pipeline is packed to be used in the queuing system, and the results are instantly displayed for the user in the form of comprehensive tables and diagrams.

For calculating off-targets we use Cas-OFFinder software, which is proven to work much more efficiently on GPUs. Unfortunately, we did not manage to obtain a dedicated webserver with powerful graphic cards, therefore we decided to increase the number of CPU cores available for calculations. If our server is overloaded with work, we will further increase the number of available cores.

We designed a consistent API for our webserver, but after multiple complex tests that took up to several days to complete, we decided to remove the possibility to submit query through API, to prevent overuse of our server. At the moment only check status and download results (in JSON format) options are available.

### Future Plans

We plan to further develop our tool. First, we are going to further expand the list of available reference genomes. We are also going to include more Cas9 variants, recognizing various PAM sequences. This will enable more comprehensive design of sgRNAs targeting specific motifs. We welcome all suggestions for improvement and development from the users via the contact details provided on the webpage.

## MATERIALS AND METHODS

### Data

Genomic coordinates for coding exons, non-coding exons, introns and transcription start sites (TSS) were downloaded from UCSC using a database interface. Full download command: mysql {genome_name} -h genome-mysql.soe.ucsc.edu -u genome -A -e ‘select * from ncbiRefSeqCurated’ -NB > {genome_file}. These data were further automatically processed to create specialized .bed files with genes and localization elements for each of available genomes. This approach is dedicated for further development of this tool to include more genomes. Whole genome sequences were downloaded as FASTA files.

### Implementation

TransCRISPR is created using Django (with Python programming language). MariaDB is used as a database, Celery with Redis as a query system, Daphne for websocket communication, Nginx as a web server, and Bootstrap-based Gentelella for a layout with Highcharts as a data visualization library. Docker with Docker Compose is used for management purposes. Off-targets are calculated using the Cas-OFFinder^25^ with a maximum of 4 (standard option) or 3 (rapid option) mismatches and later the CFD score is calculated for each off-target, as well as a cumulative CFD score^16, 25^. For on-target value calculation a dockerized version of Azimuth is used^16^.

For analyses two queue systems are available: for short calculations and for larger queries. This distinction is important because complex analyses can take up to several days, and it would be undesirable to block tasks that may take a few minutes. Where possible, parallelization and other optimizations are used to minimize time required for calculation. Software is freely available online: https://transcrispr.igcz.poznan.pl. All results are kept for 7 days. If an email address is given, an analysis completion message is sent.

Search of the motif positions is performed either with exact search in case of motifs defined as sequences with no IUPAC code or with regular expressions in case of IUPAC code in the sequence. In case of motif matrices, they are converted to IUPAC sequence using a selected rule set and then searched with regular expressions. For each sequence, reverse complementary sequence is generated and used for search on the reverse strand. For each of the found motif positions, potential guides are generated using rules for either Cas9 or dCas9.

In case of target sequences defined as coordinates, respective sequences are selected from downloaded genome files. For guide search sequences extended by 30 nucleotides before and 30 nucleotides after defined coordinates are also selected. This approach is not possible in case of sequences defined as raw sequences or in FASTA format.

Localization of DNA motifs in relation to genes is determined as follows. Firstly data downloaded from UCSC are sorted and saved to special .bed files containing data from a single chromosome, sorted by given sequence start. This operation is done only once for each genome. Found motifs position on each chromosome is also sorted. Then for each motif position on a given chromosome respective .bed file is being searched. To avoid reading the file multiple times - special mechanism is being used, where firstly there is search for the first localization from .bed file that is overlapping with the motif position. Then in a loop next positions are being read as long as part of them is overlapping with motif position. These data are also added to a buffer that is dedicated to keep them for the next motif position, as localization from .bed files may overlap multiple motifs. In case a motif is on a boundary (e.g. intron-exon, exon-intergenic etc.) or is localized in a position where different transcript variants differ with respect to intron/exon, the following hierarchy is applied: coding exon - non-coding exon - intron - intergenic.

For determination of the closest up- and downstream TSS, data from UCSC are similarly downloaded, sorted and saved to .bed files containing gene localization. During the analysis the TSSs for the closest upstream and downstream gene for each of the sorted motifs are selected and saved.

For each motif and guide, a name is generated. If a target sequence name is given (header in FASTA format or last column in .bed file) this name is used as prefix, in other cases a generic prefix is created.

### Cell culture

K562 cells were cultured in RPMI medium (Lonza, Basel, Switzerland). HEK293T, used for virus production, were cultured in DMEM medium (Lonza). Media were supplemented with 10% fetal bovine serum (Sigma-Aldrich, Saint Louis, MO, US), 2mM L-glutamine and 1% penicillin/streptomycin (Biowest, Nuaillé, France). Cells were cultured in incubator under conditions: 5% CO_2_, 37°C.

### Transduction

sgRNAs targeting selected MYC binding motifs (E-boxes) were designed using transCRISPR and cloned into lentiCRISPR_v2 vector (Addgene #52961^26^) (Table 2). Lentiviral particles were produced in HEK293T cells using 2^nd^ generation packing plasmids and Calcium Phosphate Transfection Kit (Invitrogen, Carlsbad, CA, US). K562 cells were transduced as described previously^27^. Next day 3 μg/ml of puromycin was added and cells were selected for 96 h. After selection, cells were cultured in RPMI medium with 1 μg/ml of puromycin and collected for DNA and RNA isolation 7 days post transduction.

**Table 2.**
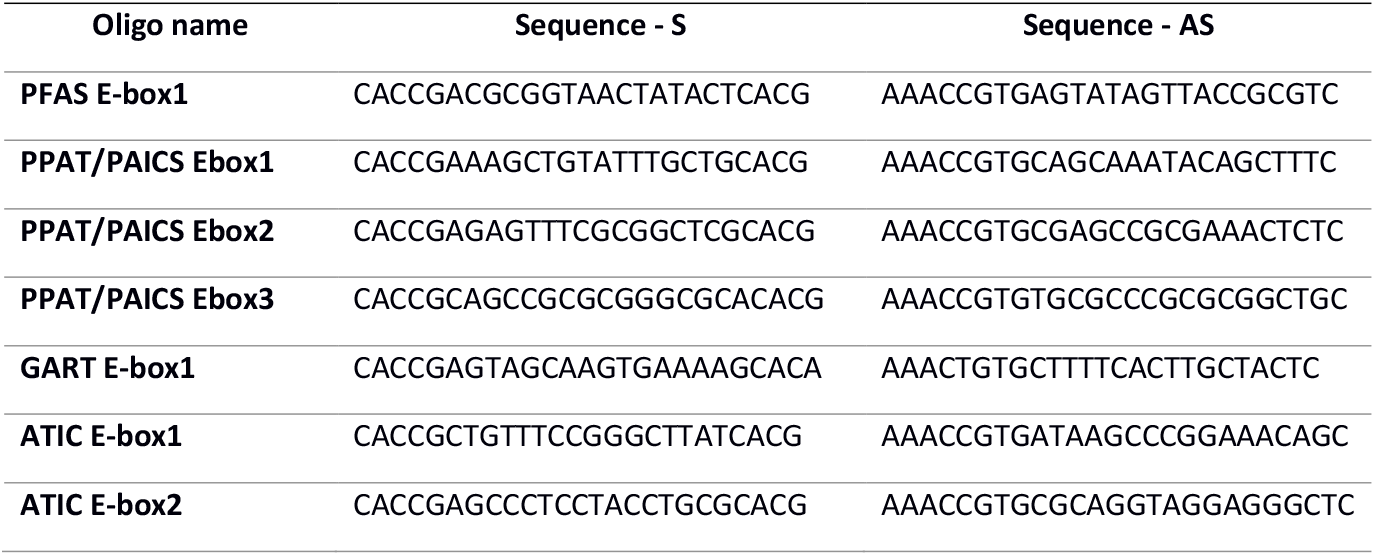
Oligo sequences.

### Cutting efficiency analysis

DNA was isolated from transduced K562 cells using Gentra Puregene Cell Kit (Qiagen, Hilden, Germany), according to the protocol. 500-800 nt DNA fragments containing regions targeted by selected sgRNAs were amplified by PCR (primers shown in Table 3). Amplicons were subjected to Sanger sequencing (Genomed, Warsaw, Poland). Chromatograms from mutated and wild type samples were compared and analyzed with TIDE calculator (https://tide-calculator.nki.nl)^28^ using indel size range of 50.

**Table 3.**
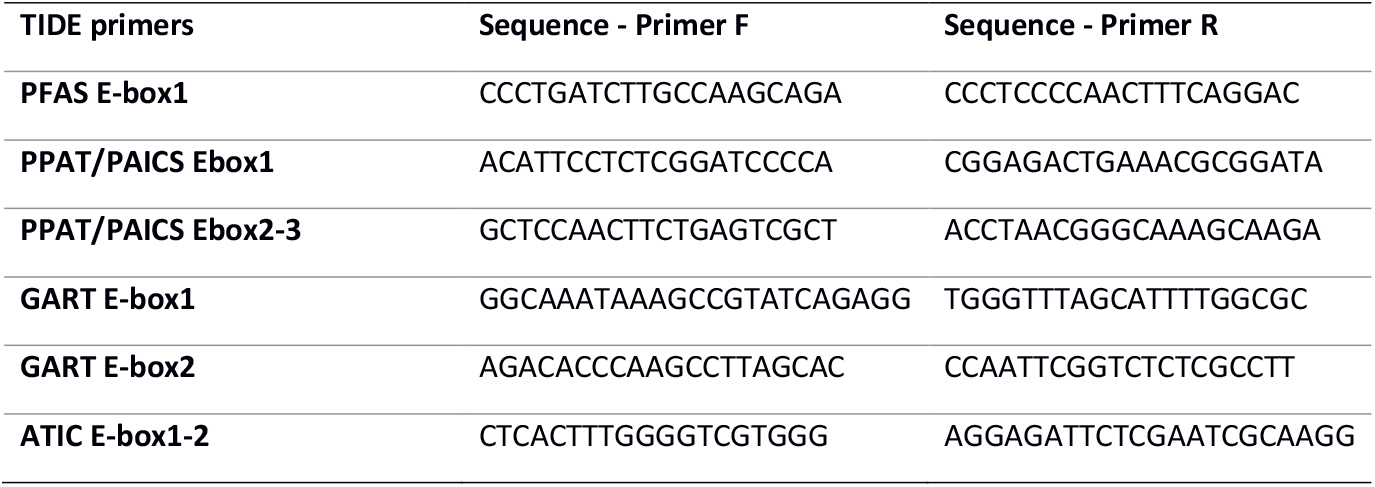
Primer sequences for TIDE analysis.

### RT-qPCR

Total RNA was isolated from transduced K562 cells using Quick-RNA^TM^ Miniprep Kit (Zymo Research, Irvine, CA, US) according to the manufacturer’s protocol. Next, 500 ng of RNA was used for cDNA synthesis using QuantiTect®Reverse Transcription Kit (Qiagen) according to the protocol. qPCR was performed with 5 ng of cDNA, PowerUp SYBR Green Master Mix (Applied Biosystems, Waltham, MA US) and primers presented in Table 4. Expression of analyzed genes was normalized to *TBP* as a reference gene. All qPCR experiments were conducted in two independent biological replicates, each with three technical replicates.

**Table 4.**
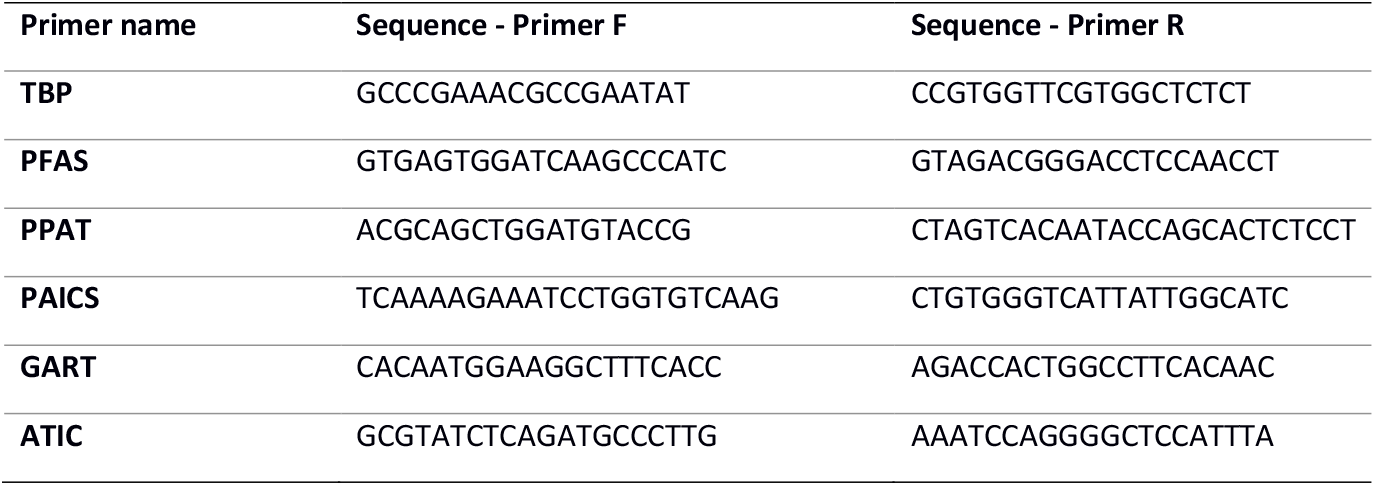
qPCR primer sequences.

## DATA AVAILABILITY STATEMENT

TransCRISPR is freely available at https://transcrispr.igcz.poznan.pl. Selected code snippets have been deposited on a github repository: https://github.com/tomaszwozniakihg/transcrispr_snippets

## AUTHORS’ CONTRIBUTIONS

TW wrote the analysis source code and website. WS, MK, MEK, MP and ADK performed tests. MK performed experimental validation. WS and MK wrote the manuscript and prepared figures. MEK and MP wrote the manual and prepared figures. ADK conceived and supervised the study, and revised the manuscript. All authors read and approved the final manuscript.

## ACKNOWLEDGEMENTS

AD-K was supported by the European Union’s Horizon 2020 research and innovation programme under grant agreement No 952304.

