## Supplementary Figure 1 for "TransCRISPR - sgRNA design tool for CRISPR/Cas9 experiments targeting DNA sequence motifs"

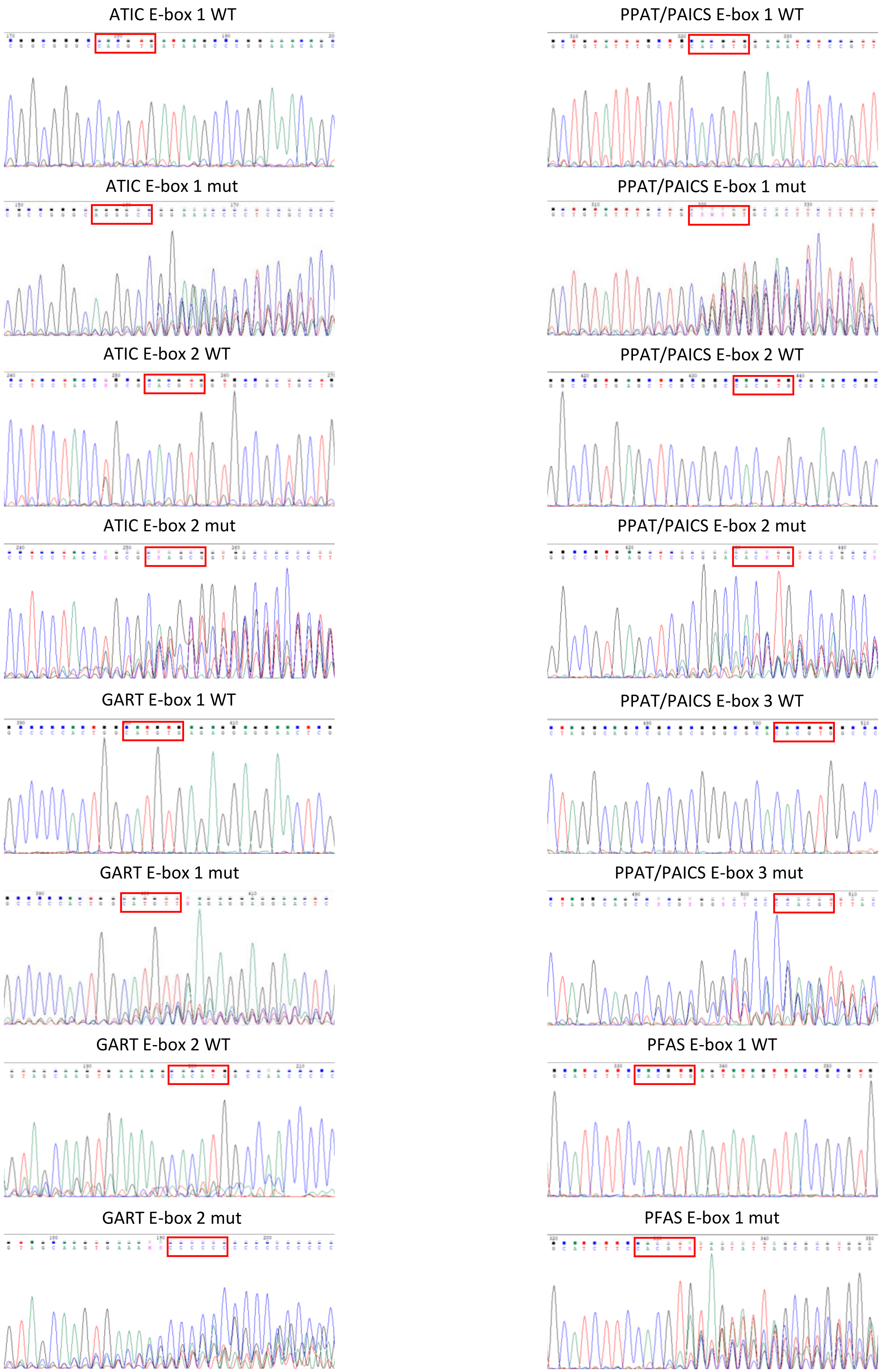


**Supplementary Figure 1. Chromatograms from Sanger sequencing of wild type and CRISPR-edited samples.**
